# Developing a model system to sustain ex vivo chloroplast function

**DOI:** 10.1101/2025.01.03.631265

**Authors:** Mariam Mohagheghi, Ali Navid, Thomas Mossington, Congwang Ye, Matthew Coleman, Steven Honag-Phou

## Abstract

Chloroplasts are critical organelles in plants and algae responsible for accumulating biomass through photosynthetic carbon fixation and cellular maintenance through metabolism in the cell. Chloroplasts are increasingly appreciated for their role in biomanufacturing, as they can produce many useful molecules, and a deeper understanding of chloroplast regulation and function would provide more insight for the biotechnological applications of these organelles. However, traditional genetic approaches to manipulate chloroplasts are slow, and generation of transgenic organisms to study their function can take weeks to months, significantly delaying the pace of research. To develop chloroplasts themselves as a quicker and more defined platform, we isolated chloroplasts from the green algae, *Chlamydomonas reinhardtii*, and examined their photosynthetic function after extraction. Combined with a metabolic modeling approach using flux-balance analysis, we identified key metabolic reactions essential to chloroplast function and leveraged this information into reagents that can be used in a “chloroplast media” capable of maintaining chloroplast photosynthetic function over time *ex vivo* compared to buffer alone. We envision this could serve as a model platform to enable more rapid design-build-test-learn cycles to study and improve chloroplast function and potentially as a foundation for the bottom-up design of a synthetic organelle-containing cell.

## 1 Introduction

Chloroplasts are critical organelles in plant and algal cells, playing key roles in metabolism and cellular maintenance. Beyond their well-known function in carbon fixation through photosynthesis, chloroplasts are increasingly recognized for their involvement in diverse metabolic pathways, such as lipid biosynthesis^1^ and amino acid production^2, 3^. These pathways make chloroplasts valuable not only for agriculture, where they can enhance crop yield^4^, but also for biomanufacturing, as they have been used to produce biofuels^5^, pharmaceuticals^6^, and recover high-value compounds such as vitamins and pigments^7-9^.

Despite their importance, studying chloroplast function through genetic approaches is laborious and time-consuming, often requiring many weeks to generate mutants even within relatively simple and well-defined model organisms such as the single celled green algae *Chlamydomonas reinhardtii*^10^. Additionally, dissecting specific biochemical pathways within chloroplasts can be challenging due to the complexity and “noise” of the cellular environment. These limitations hinder progress in both basic research and the development of biotechnological applications related to chloroplasts.

In vitro systems potentially offer a more controlled approach to study chloroplast function, removing the interference from other cellular processes. Maintaining chloroplast functionality outside of their native cellular context is difficult though, as they rapidly lose photosynthetic activity after isolation over the course of a few days^11^. However, there is evidence that they can be maintained in a prolonged viable state in non-native contexts. Notably, various sacoglossan sea slugs have demonstrated the ability to “steal” chloroplasts from their algal food sources, termed kleptoplasty, and maintain them in a photosynthetically active state for weeks to months inside their own cells^12^. Intrigued by this, we posited that there must exist a set of conditions under which chloroplasts can artificially be kept photosynthetically functional outside of their host cell.

Here, we leveraged the model organism *C. reinhardtii* and computational modeling to simulate key metabolic networks within chloroplasts and develop an initial defined chemical medium for maintaining chloroplast function *ex vivo*. We performed flux balance analysis (FBA) using a system-scale model of the *C. reinhardtii* chloroplast^13^ to analyze single-reaction knockout phenotypes. Through these simulations, we identified essential reactions and metabolites spanning several classes of molecules, notably amino acids, nucleotides, and magnesium ions. Noting the importance of protein synthesis in chloroplast function^14^, we expanded the medium to involve components commonly used *in vitro* transcription/translation buffers and showed improved maintenance of photosynthetic function of chloroplasts *ex vivo* compared to storage in sorbitol cushioned HEPES buffer alone.

## 2 Results

### 2.1 Photosynthetic function of extracted chloroplast is temperature dependent

To determine whether functional chloroplast lifetimes can be modulated outside of their host cell, we extracted chloroplasts from *C. reinhardtii* and measured their photosynthetic function after extraction using pulse amplitude modulated (PAM) fluorometry (Fig. 1A). Percoll/ficoll density gradients allowed for the efficient separation of intact chloroplasts from whole cells and thylakoid membranes^15^ (Fig 1B) where strong membrane-bound chlorophyll α signal could be detected in individual chloroplasts. Because *C. reinhardtii* contain only a single chloroplast per cell, we were able to measure our recovery rate by counting the number of isolated chloroplasts, obtaining >70% recovery on average (Fig. 1C).

**Figure 1.**
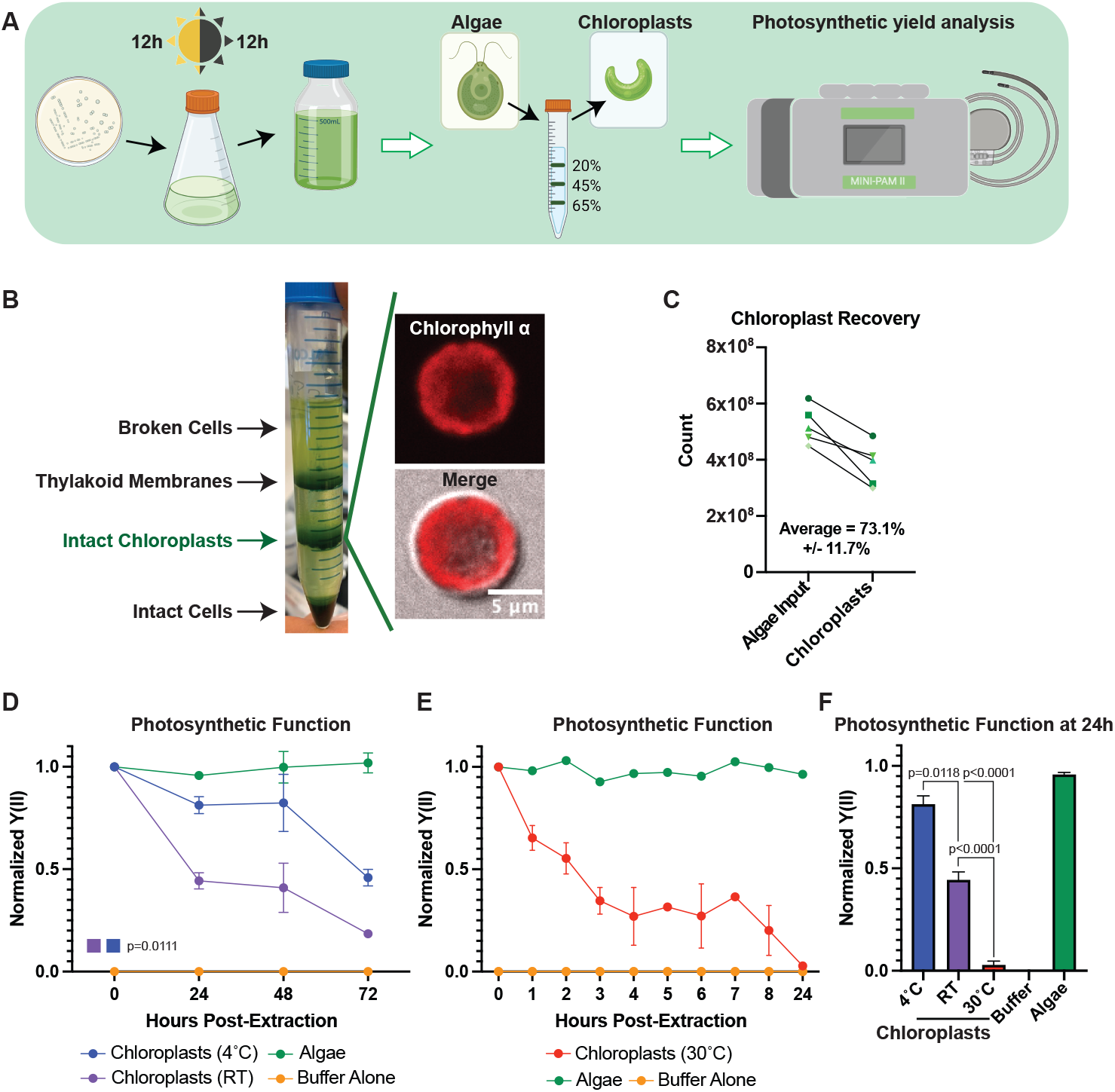
Chloroplast function ex vivo can be modulated by temperature. **A)** Schematic describing the experimental pipeline for *C. reinhardtii* culture, chloroplast extraction, and photosynthetic measurements. **B)** Image of representative chloroplast extraction with (inset) confocal micrographs of an extracted chloroplast. **C)** Line graph showing chloroplast recovery as a percent of starting algal cells. **D)** Line graphs showing normalized Y(II) values over time at 4°C and RT. P-values calculated using repeated measures 2-way ANOVA. **E)** Line graphs showing normalized Y(II) values over time at 30°C. **F)** Bar graph comparing normalized Y(II) values at 24h post extraction at various temperatures. P-values calculated by Student’s t-test.

Chloroplasts have a limited functional lifespan after extraction from their host cell. Indeed, we observed results consistent with those reported in the literature^11^, where most chloroplast photosynthetic functionality was lost within 72 hours of extraction in sorbitol cushioned HEPES buffer (HEPES buffer) (Fig. 1D). We observed a sharp decline in the effective quantum yield of photosystem II (Y(II)) through PAM measurements within the first 24 hours, before tapering off through 72 hours, when stored at room temperature (RT). We observed a significant delay in the rate of Y(II) decline in extracted chloroplasts when stored at 4°C compared to RT (p=0.0111), suggesting that chloroplast viability after removal from host cells could be influenced by temperature. Increasing the incubation temperature of extracted chloroplasts to 30°C accelerated the rate of Y(II) decline, demonstrating a similar loss of photosynthetic function within 8 hours that is typically seen in 72 hours at RT (Fig. 1E), with the differences sharply highlighted at 24h (Fig. 1F), and we reasoned this could serve as a useful assay for measuring chloroplast function moving forward.

### 2.2 Computational analysis of *C. Reinhardtii* identifies metabolites essential to chloroplast function

Encouraged by our initial results, we sought to identify other conditions that could enhance metabolic robustness of chloroplasts or extend functional lifetimes when decoupled from their host cells to develop a chloroplast “growth media”. We performed computational system-level examination of chloroplasts using flux balance analysis (FBA) on a published genome-scale model (GEM) of algal chloroplasts from three algal species^16^. GEMs use an organism’s annotated genome to generate a mathematical reconstruction of their metabolic networks in a stoichiometric matrix that can be probed to identify characteristics such as important pathways for growth, robustness to genetic and environmental perturbations, and tradeoffs between different system objectives^17^. While the published GEM contained reactions from multiple algal species, we used only the reactions related to *C. reinhardtii* for our analyses.

To identify essential reactions, we performed systematic *in silico* single reaction knockouts of each metabolic reaction in the model, specifically focusing on identifying sets of reactions that are essential for growth or ATP generation in the chloroplast (Fig. 2A). Our initial results included both internal chloroplast biochemical reactions as well as transport processes that facilitate import of nutrients and export of waste material (Supplementary Table 1). To generate a list of essential *C. reinhartii* cytosol metabolites that need to be imported for operation of its chloroplast, we excluded the list of essential internal biochemical reactions and focused on the essential transport reactions. Specifically, we looked at import, reasoning that any nutrients requiring import to chloroplasts from the host cell would also be required in a defined medium to support chloroplast maintenance and operation and obtained a list of imported metabolites.

**Figure 2.**
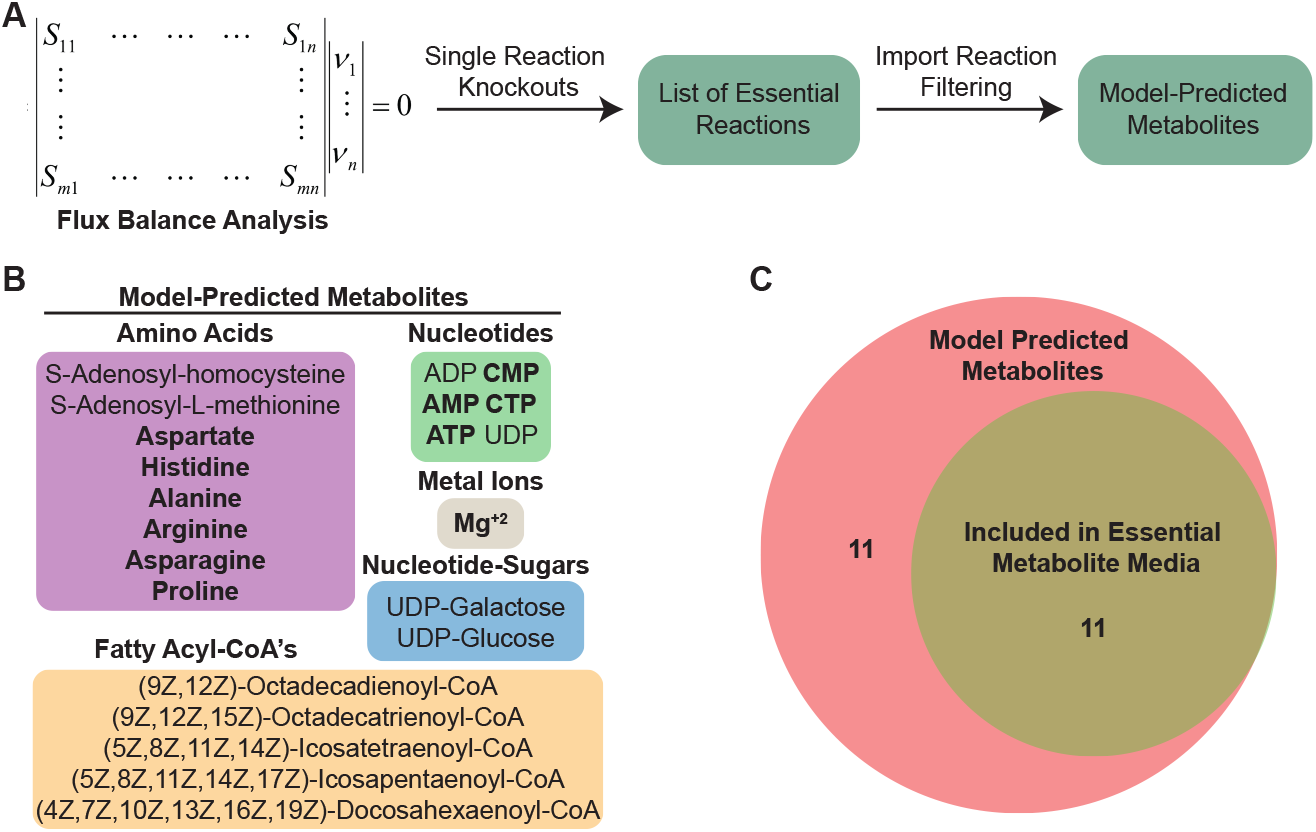
Flux balance analysis predicted a set of essential imported metabolites. **A)** Schematic outlining the computational pipeline used to determine essential imported metabolites in *C. reinhardtii* chloroplasts. **B)** List of essential imported metabolites predicts from *C. reinhardtii* chloroplast model. **C)** Venn diagram showing number of metabolites included in EM media.

Among the model-predicted metabolites are various amino acids, fatty acid chains, nucleotides, nucleotide-sugars, cofactors, and critical biosynthesis intermediates (Fig. 2B). In general, these compounds are necessary for DNA replication and protein production, maintaining chloroplast membranes^18,19^, and generating defensive molecules such as antioxidants that are needed to negate the deleterious effects of ROS produced by photosynthesis^20^. Since it is known that photosynthesis is an inherently damaging process resulting in high protein turnover^21^, we focused on the metabolites involved in DNA and protein synthesis, and excluded the components more related to supporting membrane integrity. From the model-predicted metabolite list, we chose 11 metabolites to include in an “essential metabolite chloroplast media” (EM) (Fig. 2C).

### 2.3 Computationally identified metabolites prolong chloroplast photosynthetic function over time after extraction

To determine whether the identified metabolites could augment extracted chloroplast function, we incubated chloroplasts at 30°C either in HEPES buffer or EM media and took PAM readings over time (Fig. 3A). While we saw an encouraging trend towards chloroplasts maintaining a higher photosynthetic yield (Y(II)) in the EM media, it was not significant (p=0.2894). Closer examination of the EM media revealed a prevalence of amino acids, nucleotides, and importantly magnesium ions, which are essential for transcription and translation. Due to the high demand active photosynthesis places on chloroplasts, we reasoned that any media capable of supporting chloroplast function must also provide the raw materials needed to replace damaged proteins. Thus, we decided to use the PANOx-SP reaction buffer common to transcription/translation cell-free protein synthesis (CFPS) systems^22^, which support those processes in vitro, as an “enhanced essential metabolite” (EEM) media (Supplementary Table 2). EEM media encompassed nearly all EM media components and provided additional amino acids, nucleotides, cofactors, salts, and energy rich components (Fig. 3B). Incubation in EEM media showed a significantly slower decline in photosynthetic function over time compared to incubation in HEPES buffer (Fig. 3C). To determine whether there were any morphological changes happening concurrently with photosynthetic decline, we imaged extracted chloroplasts on a confocal microscope after incubation at 30°C in HEPES buffer, EM, or EEM media. We categorized chloroplast morphology based on their shape and membrane appearance as either “intact”, “intermediate”, or “degraded” (Fig. 3D). We were able to distinguish all three categories consistently between the different buffer conditions and quantified the population percentage of each category at 0h and 8h of incubation. The percentages between each group were consistent at the initial timepoint, however we saw differences emerge at 8h (Fig. 3E). Although both EM and EEM media had similar amounts of intact chloroplasts at 8h, both groups performed better than HEPES buffer. Interestingly, we counted less degraded chloroplasts in EEM media compared to EM media (p=0.0152) and HEPES buffer (p=0.0026) at 8h.

**Figure 3.**
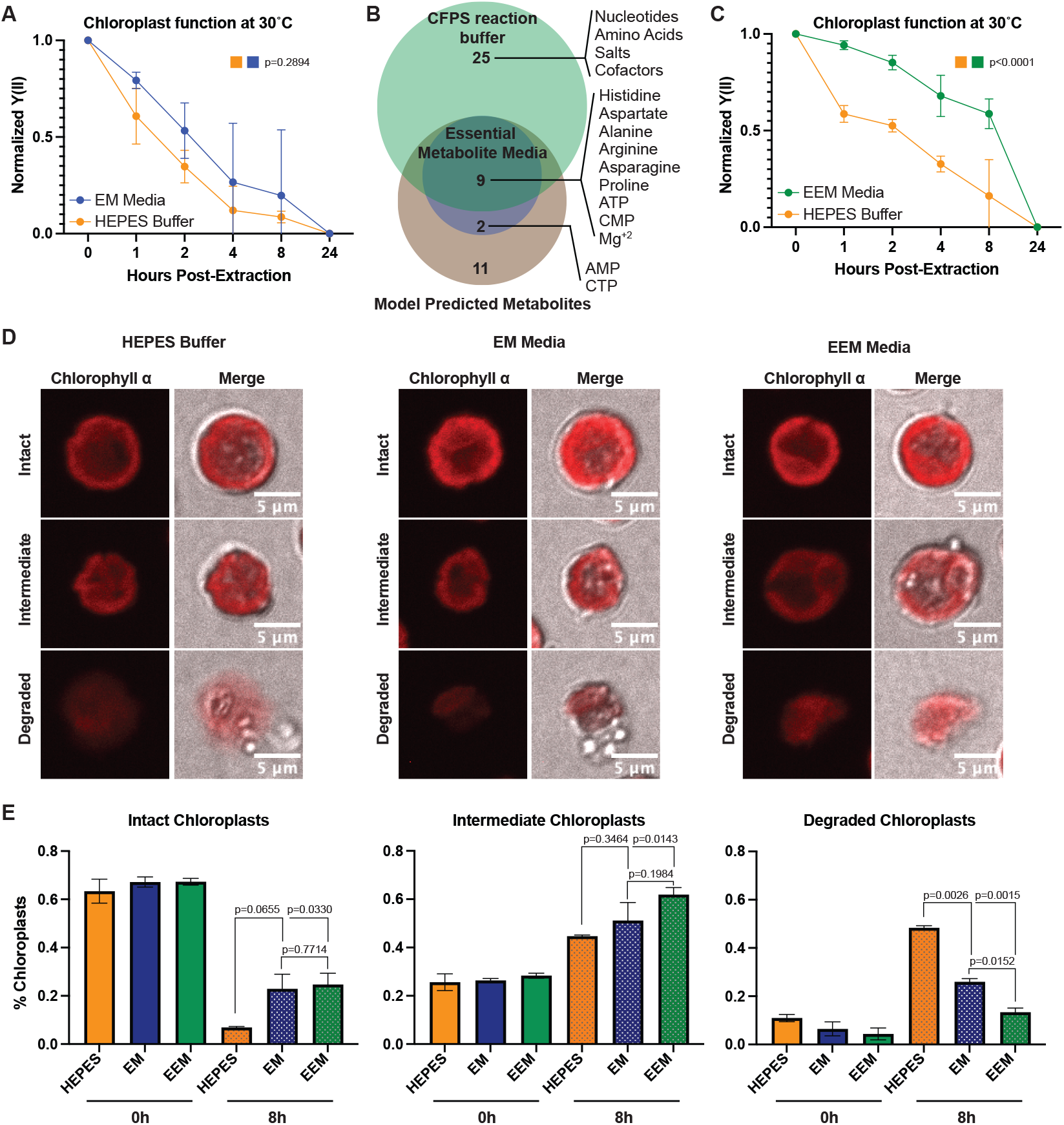
Metabolites supporting transcription and translation prolong extracted chloroplast photosynthetic function over time. **A)** Line graph showing normalized Y(II) values over time from extracted chloroplasts at 30°C in HEPES buffer and EM media. N=3 experiments. P-values calculated using repeated measures 2-way ANOVA. **B)** Venn diagram showing overlap between predicted metabolites, EM media, and CFPS reaction buffer components. **C)** Line graph showing normalized Y(II) values over time from extracted chloroplasts at 30°C in HEPES buffer and EEM media. N=4 experiments. P-values calculated using repeated measures 2-way ANOVA. **D)** Confocal micrographs showing representative intact, intermediate, and degraded chloroplasts in HEPES, EM media, and EEM media. **E)** Bar graphs showing quantification of intact, intermediate, and degraded chloroplasts after incubation at 30°C for 0h or 8h. N=2 experiments; 4 fields of view per group per experiment. P-values calculated by Student’s t-test.

## 3 Discussion

The world population is expected to grow beyond nine billion by 2050^23^, increasing demands on food production and energy infrastructure. Improvements to existing crop yields and alternative energy sources utilizing bioenergy are prospective solutions to these challenges. Central to both processes are chloroplasts, which are mature plastids found in plants and algae that perform carbon fixation through photosynthesis, among other metabolic processes. Chloroplasts have become increasingly appreciated for their potential as biotechnology workhorses, proving instrumental in producing biomolecules for therapeutics^24^ and lipids used in biofuel production^25,26^. However, traditional methods of studying and manipulating chloroplasts within cells rely on labor-intensive techniques, often requiring many generations to achieve homoplasy, significantly slowing down research progress.

Our study sought to approach this issue from a different angle and take the initial steps towards developing an *ex vivo* chloroplast platform. Working with chloroplasts outside their cellular environment could simplify studies, eliminating interference from other cellular processes and provide a more controlled system. However, chloroplasts rapidly lose photosynthetic function after extraction from their host cells. We took inspiration from Sacoglassan sea slug kleptoplasty, which can maintain photosynthetically active chloroplasts from their algal sources for months after consumption^12^. In fact, many species of unicellular organisms have displayed the ability to maintain active chloroplasts in completely non-native contexts (reviewed in ^27^), and we deduced that there must exist some external set of conditions that could satisfy the requirement of keeping them viable.

Here, we successfully isolated chloroplasts from *C. reinhardtii* and identified a set of metabolites that could slow the decline of photosynthetic function after extraction. We used a *C. reinhardtii* chloroplast metabolic model and performed flux balance analysis (FBA) across the known metabolic reactions present in the model. Modeling single-reaction knockouts and optimizing for ATP production and biomass accumulation in the chloroplast, we identified a list of metabolic reactions and essential metabolites imported into chloroplasts and developed our initial media recipe. While our initial media (EM) did not significantly maintain chloroplast function at 30°C, we were encouraged by the findings. We noticed an abundance of transcription and translation related components in EM media, which led us to infer several key points: 1) These components are crucial since photosynthesis itself damages the photosynthetic machinery within chloroplasts^28-30^; 2) supporting protein synthesis and turnover is critical for maintaining function; and 3) CFPS reaction buffers could satisfy this requirement. CFPS enables protein production using cell extracts, and typical *E. coli* PANOx-SP^22^ CFPS buffers contain most of the components in EM media, along with additional amino acids, nucleotides, salts, and cofactors. When we incubated extracted chloroplasts in EEM media at 30°C over time, we observed significant delays in chloroplast functional decline through PAM measurements. Together with our morphological analyses, our data showed that chloroplasts in EEM media were less degraded and maintained better membrane integrity, suggesting that EEM media better supported their structure and function over time.

However, like any computational approach, our results depend on the accuracy of the underlying metabolic model. Our analysis identified several additional components absent in both the EM and EEM media, including adenosyl-containing amino acids, UDP-linked sugars, and fatty acid cofactors. S-adenosyl-homocysteine (AdoCys) can be converted into S-adenosyl-L-methionine (AdoMet), which serves as a universal methyl donor for methylation reactions and is a precursor for polyamine synthesis, which are critical for cell growth, division, and lipid accumulation^31,32^. Although AdoCys and AdoMet are absent in the media, downstream polyamines like putrescine and spermidine are present in the EEM media, potentially compensating for this absence. External polyamines, such as putrescsine and spermidine, are known to influence *C. Reinhardtii* growth and may be taken up in limited amounts^33,34^, though whether extracted chloroplasts retain these uptake mechanisms remains unexplored.

UDP-glucose, UDP-galactose, and the long- and very long-chain fatty acyl CoA’s we identified likely play important roles in lipid production, membrane maintenance, and fatty acid metabolism^35,36^. UDP-glucose and UDP-galactose are interconvertible, with UDP-galactose serving as a key component of two photosynthetic membrane galactolipids: Monogalactosyldiacylglycerol (MGDG) and digalactosyldiacylglycerol (DGDG)^18,37^. These lipids are highly conserved across photosynthetic organisms and can be produced entirely within chloroplasts via the “prokaryotic pathway” or in cooperation with the endoplasmic reticulum (ER) through the “eukaryotic pathway” which involves lipid trafficking between the chloroplast and ER. Both MGDG and DGDG serve critical structural and functional roles in thylakoid membranes and their associated photosystem complexes^38^. Although extracted chloroplasts in EEM media maintained higher photosynthetic yield than the other conditions, we still observed a noticeable increase in the amount of degraded or membrane-damaged chloroplasts by 8h and a sharp drop-off in photosynthetic yield by 24h at 30°C. Although we did not explore the role of these lipid components, these data would suggest that the chloroplasts may not have been sufficiently able to maintain their lipid membranes after extraction, contributing to their loss of function, and we speculate that inclusion of these components could potentially further prolong extracted chloroplast function over time.

In this study, incubation of extracted chloroplasts in EEM media significantly delayed the decline of photosynthetic function compared to HEPES buffer and EM media. While the exact mechanism by which EEM media prolongs chloroplast function is unknown, we speculate the essential components are necessary to support transcription, translation, and protein turnover within the chloroplast. Identifying media formulations that can sustain chloroplast functionality ex vivo for extended periods could facilitate their direct manipulation and create a more efficient research and biomanufacturing^39^ platform. Refining the chloroplast media formulation by incorporating additional metabolites, such as fatty acid cofactors and UDP-linked sugars, could clarify their roles in maintaining chloroplast function. These advances could potentially lead to faster design-build-test-learn cycles for chloroplast research and biotechnological applications. Additionally, a deeper understanding of the conditions required to maintain critical organelles like chloroplasts outside of their host cells may advance efforts in constructing synthetic organelle-containing cells from the bottom up.

## 4 Materials and Methods

### Strains and Growth Conditions

*Chlamydomonas reinhardtii* strain CC400 cw15 mt+ was acquired from the Chlamydomonas Resource Center. *C. reinhardtii* was cultured in 250mL flasks using high salt with acetate (HSA) media supplemented with Hutner’s trace elements (Chlamydomonas Resource Center) under 12h:12h light/dark cycles. Cultures were expanded to 500mL bottles for chloroplast extractions under the same growth conditions and harvested after 4-5 days.

### Chloroplast extraction

Chloroplasts were extracted from *C. Reinhardtii* cells according to *Mason et al*.^15^ with slight modifications. Briefly, 5×10^6^-1×10^7^ cells were harvested from a 500ml culture by centrifugation at 4000g for 10 minutes at 4°C, washed with 10mL HEPES buffer (50mM HEPES-KOH pH 7.5), pelleted, and resuspended in 2mL HEPES buffer. The algae were diluted by adding 8-10mL of isolation buffer (50mM HEPES-KOH pH 7.5, 2mM EDTA, 1mM MgCl_2_, 300mM Sorbitol, and 1% BSA), and lysed by a single pass through a 27-gauge needle. Lysed algae were centrifuged at 750g for 2 minutes at 4°C and gently resuspended in 2mL of isolation buffer using a paintbrush before layering on top of a pre-prepared Percoll/Ficoll gradient without isoascorbate. Gradients were centrifuged at 3265g for 15 minutes at 4°C and the band at the 45%-65% interface was collected as intact chloroplasts before dilution with 50mM HEPES-KOH pH 7.5 and used for downstream assays.

### Measurement of photosynthetic efficiency (MINI PAM-II)

The photochemical activity of Photosystem II (PSII) was measured using a pulse-amplitude modulated (PAM) fluorometer (MINI-PAM-II, Heinz-Walz, Germany). Chloroplast concentrations were determined via hemocytometer and aliquoted into sterile Eppendorf tubes at a concentration of 1×10^6^ chloroplasts/mL. Chloroplasts were pelleted at 600g for 3 minutes at room temperature and resuspended in equal volumes of HEPES buffer, EM, or EEM media to maintain consistent concentrations across samples. The centrifugation and resuspension processes were repeated to eliminate residual extraction and storage buffers. Resuspended samples were incubated at pre-determined temperatures (4°C, 25°C, and 30°C) and protected from light with foil. Time “0” chloroplast measurements were taken immediately upon resuspension into designated buffer post-extraction. 200µl aliquots of 1×10^6^ chloroplasts/mL stock solution were pipetted into a MINI PAM-II chamber and read per manufacturer instructions. N=3 experiments and n=4 experiments were performed to compare HEPES buffer to EM media and HEPES buffer to EEM media, respectively. Repeated measures two-way ANOVA with Geisser-Greenhouse correction was used to determine statistical significance of changes in Y(II) values over time.

### Microscopy and Image Analysis

Extracted chloroplasts were pelleted and gently resuspended in HEPES buffer, EM, or EEM media at a concentration of 1×10^6^ chloroplasts/mL. 15µl of the suspension was plated onto a No. 1.5 18-well glass slide (Ibidi #81821) and allowed to settle for 30 minutes. Differential interference contrast (DIC) and fluorescence (415nm ex/640nm em) images were captured using a 60x objective on a Nikon inverted confocal microscope. Morphology was classified into three categories: Intact (fully intact membrane with minimal degradation), intermediate (some visible membrane perforation), and degraded (no intact membrane). For each timepoint (n=2 experiments per timepoint), 4 images containing 250-500 chloroplasts each were analyzed using ImageJ (ImageJ2 2.14.0/1.54f) and classifications were averaged across the images. Statistical analysis was performed using Student’s t-test to assess significance.

### Flux Balance Analysis

Flux balance analysis (FBA) is a constraint-based reconstruction and analysis approach that uses a genome-scale metabolic reconstruction as its basis. The elementary functional information derived from annotated genomes is used with available knowledge of enzymology to reconstruct all metabolic pathways in an organism. Metabolism is represented mathematically as a stoichiometric matrix, *S(m×n)*, where *m* is the number of metabolites and *n* the number of different reactions. By assuming mass balance and the system is in metabolic steady-state, the following set of linear equations govern the system’s behavior:

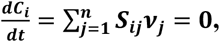

where *C*_*i*_ is the concentration of metabolite *i*. Other limitations are imposed on the system based on experimental studies, such as bounding the flux through a reaction, as well as constraints on the import of nutrients and export of waste products. These constraints are formulated as:

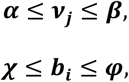

where *b*_*i*_ and ν_*j*_ are the export/import flux of metabolite species *i*, and the flux through internal reaction *j* respectively. *α, β, χ*, and *φ* are the lower and upper limits for these fluxes. Finally, FBA employs linear programming to determine a feasible steady-state flux vector that optimizes an objective function, often chosen to be the production of biomass, *i*.*e*. cellular growth. For our analyses, FBA was used with a published system-scale model of algal chloroplasts^40^ to analyze single gene and reaction knockout phenotypes in the model. While we only used the *C. reinhardtii*-related reactions in the model, since the developers of the system-scale model used *Nannochloropsis gaditana* as a base for all models, any reaction in the *C. reinhardtii* model that is also present in *N. gaditana* has a “Nano” designation.

For the single reaction knockouts, the activity of each reaction in the model is blocked and the model is tested for its ability to grow (i.e., produce biomass). When blocking the activity of a reaction leads to cessation of growth, that reaction is labeled an essential reaction for the analyzed environment. In case of single gene knockout, for inactivation of each gene, the reactions that are catalyzed by the product of translation of that protein are blocked and if this results in cessation of growth, the gene is deemed essential.

For our analysis identifying essential media nutrients, the list of essential transport reactions, particularly import reactions were used to predict an essential medium for stand-alone operation of *C. reinhardtii* chloroplast. The transport reactions in geome-scale models of metabolism typically are included in the models based on annotation of membrane surface proteins as well as known auxotrophies of a system. In some cases, when a biosynthetic pathway for a known component of biomass is missing in the metabolic reconstruction, the trasnsport reactions are “gap-filled” into the models to ensure its ability to produce biomass.

