## Supplemental Table 2 for "Developing a model system to sustain ex vivo chloroplast function"

**Supplementary table 2. Enhanced metabolite media formulation**

| | **Concentration** |
|---|---|
| **ATP** | 1.2mM |
| **GMP** | 0.86mM |
| **UMP** | 0.86mM |
| **CMP** | 0.86mM |
| **E. coli tRNA** | 170ug/mL |
| **19 Essential Amino Acids (excluding glutamate)** | 2mM |
| **NAD** | 0.33mM |
| **Coenzyme A** | 0.27mM |
| **Spermidine** | 1.5mM |
| **Putrescine** | 1mM |
| **Potassium oxalate** | 2.7mM |
| **Potassium glutamate** | 175mM |
| **Ammonium glutamate** | 10mM |
| **Magnesium glutamate** | 10mM |
| **PEP** | 33mM |
