## Supplemental Table 1 for "Developing a model system to sustain ex vivo chloroplast function"

**Supplementary table 1. List of essential reactions in chloroplasts**

| Reaction ID | Chloroplast Essential Reactions |
|---|---|
| NanoG0117 | 'homoserine dehydrogenase' |
| NanoG0124 | threonine synthase' |
| NanoG0130 | 'atp:l-homoserine o-phosphotransferase' |
| NanoG0131 | 'threonine ammonia-lyase' |
| NanoG0189 | 'aspartate---prephenate aminotransferase' |
| NanoG0191 | 'Chorismate synthase' |
| NanoG0193 | '3-phosphoshikimate 1-carboxyvinyltransferase' |
| NanoG0199 | 'Phosphoribosylanthranilate isomerase' |
| NanoG0202 | '3-dehydroquinate synthase' |
| NanoG0203 | 'shikimate kinase' |
| NanoG0204 | 'quinate/shikimate dehydrogenase' |
| NanoG0205 | 'anthranilate phosphoribosyltransferase' |
| NanoG0209 | 'anthranilate synthase' |
| NanoG0210 | '3-dehydroquinate dehydratase' |
| NanoG0211 | '3-deoxy-7-phosphoheptulonate synthase' |
| NanoG0258 | 'branched-chain-amino-acid transaminase' |
| NanoG0261 | 'acetolactate synthase' |
| NanoG0263 | 'ketol-acid reductoisomerase' |
| NanoG0276 | 'dihydroxy-acid dehydratase' |
| NanoG0281 | 'branched-chain-amino-acid transaminase' |
| NanoG0293 | '4-hydroxy-tetrahydrodipicolinate synthase' |
| NanoG0324 | 'udp-sulfoquinovose synthase' |
| NanoG0348 | 'l-ascorbate peroxidase' |
| NanoG0350 | 'glutathione dehydrogenase (ascorbate)' |
| NanoG0401 | 'triose-phosphate isomerase' |
| NanoG0412 | 'phosphoglycerate kinase' |
| NanoG0514 | 'ribulose-phosphate 3-epimerase' |
| NanoG0516 | 'transketolase' |
| NanoG0528 | 'ribose-phosphate diphosphokinase' |
| NanoG0588 | 'phosphoribulokinase' |
| NanoG0589 | 'ribulose-bisphosphate carboxylase' |
| NanoG0609 | 'inorganic diphosphatase' |
| NanoG0677 | '3-hydroxyacyl-[acyl-carrier-protein] dehydratase (n-c12:0)' |
| NanoG0679 | '3-hydroxyacyl-[acyl-carrier-protein] dehydratase (n-c16:0)' |
| NanoG0681 | '3-hydroxyacyl-[acyl-carrier-protein] dehydratase' |
| NanoG0683 | '3-hydroxyacyl-[acyl-carrier-protein] dehydratase' |

| | |
|---|---|
| NanoG0685 | 'enoyl-[acyl-carrier-protein] reductase (nadph, re-specific)' |
| NanoG0688 | '3-oxoacyl-[acyl-carrier-protein] reductase [(n-c4:0)]' |
| NanoG0690 | '3-oxoacyl-[acyl-carrier-protein] reductase [(n-c6:0)]' |
| NanoG0692 | '3-oxoacyl-[acyl-carrier-protein] reductase (n-c8:0)' |
| NanoG0694 | '3-oxoacyl-[acyl-carrier-protein] reductase (n-c10:0)' |
| NanoG0696 | '3-oxoacyl-[acyl-carrier-protein] reductase (n-c12:0)' |
| NanoG0698 | '3-oxoacyl-[acyl-carrier-protein] reductase (n-c16:0)' |
| NanoG0699 | '3-oxoacyl-[acyl-carrier-protein] reductase' |
| NanoG0704 | 'malonyl-coa:[acyl-carrier-protein] s-malonyltransferase/ malonyl-coa:acp-trans-acylase' |
| NanoG0712 | '3-hydroxyacyl-[acyl-carrier-protein] dehydratase' |
| NanoG0714 | 'Acetyl transacylase' |
| NanoG0728 | 'stearoyl-[acyl-carrier-protein] delta9-desaturase ((9Z)-n-C18:1)' |
| NanoG0735 | 'enoyl-[acyl-carrier-protein] reductase (nadh)[n-c18:0]' |
| NanoG0736 | 'enoyl-[acyl-carrier-protein] reductase (nadh)' |
| NanoG0740 | 'enoyl-[acyl-carrier-protein] reductase (nadh)' |
| NanoG0742 | '3-hydroxyacyl-[acyl-carrier-protein] dehydratase (n-c8:0)' |
| NanoG0744 | '3-hydroxyacyl-[acyl-carrier-protein] dehydratase [(n-c6:0)]' |
| NanoG0746 | '3-hydroxyacyl-[acyl-carrier-protein] dehydratase (n-c18:0)' |
| NanoG0755 | 'acyl-[acyl-carrier-protein] delta9-desaturase ((9Z)-n-C16:1)' |
| NanoG0771 | 'beta-ketoacyl-[acyl-carrier-protein] synthase i' |
| NanoG0785 | '3-oxoacyl-[acyl-carrier-protein] synthase (C4:0 forming)' |
| NanoG0793 | '3-oxoacyl-[acyl-carrier-protein] reductase (n-c18:0)' |
| NanoG0802 | 'acetyl-coa carboxylase' |
| NanoG1695 | 'galactolipid galactosyltransferase' |
| NanoG0892 | 'Phosphatidate phosphatase (n-C14 0)' |
| NanoG0894 | 'Phosphatidate phosphatase (n-C16 0)' |
| NanoG0896 | 'Phosphatidate phosphatase (n-C16 1)' |
| NanoG0898 | 'Phosphatidate phosphatase (n-C18 0)' |
| NanoG0900 | 'Phosphatidate phosphatase (n-C18 1)' |
| NanoG0902 | 'Phosphatidate phosphatase (n-C18 2)' |
| NanoG0904 | 'Phosphatidate phosphatase (n-C18 3)' |
| NanoG0908 | 'Phosphatidate phosphatase (n-C20 4)' |
| NanoG0910 | 'Phosphatidate phosphatase (n-C20 5)' |
| NanoG0911 | '1-acylglycerol-3-phosphate O-acyltransferase (n-C14:0)' |
| NanoG0912 | '1-acylglycerol-3-phosphate o-acyltransferase (n-C16:1)' |
| NanoG0913 | '1-acylglycerol-3-phosphate o-acyltransferase (n-C18:0)' |
| NanoG0914 | '1-acylglycerol-3-phosphate o-acyltransferase (n-C18:1)' |
| NanoG0923 | 'sulfoquinovosyltransferase (n-C16 0)' |
| NanoG0924 | 'sulfoquinovosyltransferase (n-C16 1)' |

| | |
|---|---|
| NanoG0925 | 'sulfoquinovosyltransferase (n-C18 1)' |
| NanoG0926 | 'sulfoquinovosyltransferase (n-C18 2)' |
| NanoG0927 | 'sulfoquinovosyltransferase (n-C20 4)' |
| NanoG0928 | 'sulfoquinovosyltransferase (n-C20 5)' |
| NanoG0959 | '1-acylglycerol-3-phosphate o-acyltransferase (n-C16:0)' |
| NanoG0973 | 'UDPgalactose 1,2-diacylglycerol 3-beta-D-galactosyltransferase (n-C14 0)' |
| NanoG0975 | 'UDPgalactose 1,2-diacylglycerol 3-beta-D-galactosyltransferase (n-C16 0)' |
| NanoG0977 | 'UDPgalactose 1,2-diacylglycerol 3-beta-D-galactosyltransferase (n-C16 1)' |
| NanoG0979 | 'UDPgalactose 1,2-diacylglycerol 3-beta-D-galactosyltransferase (n-C18 1)' |
| NanoG0981 | 'UDPgalactose 1,2-diacylglycerol 3-beta-D-galactosyltransferase (n-C20 4)' |
| NanoG0983 | 'UDPgalactose 1,2-diacylglycerol 3-beta-D-galactosyltransferase (n-C20 5)' |
| NanoG0996 | 'glycerol-3-phosphate: acyl-coa acyltransferase C18:2' |
| NanoG0998 | 'glycerol-3-phosphate: acyl-coa acyltransferase C18:3' |
| NanoG1000 | 'glycerol-3-phosphate: acyl-coa acyltransferase C20:4' |
| NanoG1002 | 'glycerol-3-phosphate: acyl-coa acyltransferase C20:5' |
| NanoG1620 | 'sulfoquinovosyltransferase (n-C14 0)' |
| NanoG1631 | 'UDPgalactose 1,2-diacylglycerol 3-beta-D-galactosyltransferase (n-C18 2)' |
| NanoG1035 | 'glycerol-3-phosphate dehydrogenase [nad(p)+]' |
| NanoG1268 | 'glutamate-tRNA ligase' |
| NanoG1269 | 'Mg-protoporphyrin IX monomethyl ester (oxidative) cyclase I' |
| NanoG1270 | 'Mg-protoporphyrin IX monomethyl ester (oxidative) cyclase II' |
| NanoG1271 | 'Mg-protoporphyrin IX monomethyl ester (oxidative) cyclase III' |
| NanoG1273 | 'hydroxymethylbilane synthase' |
| NanoG1276 | 'magnesium protoporphyrin ix methyltransferase' |
| NanoG1278 | 'porphobilinogen synthase' |
| NanoG1279 | 'protochlorophyllide reductase' |
| NanoG1281 | 'chlorophyllide a oxygenase' |
| NanoG1282 | 'chlorphyll b reductase; chlorophyllide a oxygenase' |
| NanoG1283 | 'protoporphyrinogen oxidase' |
| NanoG1285 | 'glutamate-1-semialdehyde 2,1-aminomutase' |
| NanoG1289 | 'glutamyl-trna reductase' |
| NanoG1293 | 'coproporphyrinogen oxidase' |
| NanoG1294 | 'divinyl chlorophyllide a 8-vinyl-reductase' |
| NanoG1296 | 'chlorophyll synthase' |
| NanoG1302 | 'uroporphyrinogen decarboxylase' |
| NanoG1304 | 'uroporphyrinogen-iii synthase' |
| NanoG1310 | 'magnesium chelatase' |
| NanoG1338 | '1-deoxy-d-xylulose-5-phosphate synthase' |
| NanoG1383 | 'glutathione-disulfide reductase' |

| | |
|---|---|
| NanoG1397 | 'lycopene cyclase (alpha-carotene producing)' |
| NanoG1398 | 'lycopene cyclase (delta-carotene producing)' |
| NanoG1399 | 'Neurosporene oxidoreductase' |
| NanoG1400 | 'phytoene desaturase (2)' |
| NanoG1404 | 'zeta-carotene desaturase' |
| NanoG1405 | 'alpha-carotene hydroxylase (alpha-cryptoxanthin forming)' |
| NanoG1408 | 'lycopene beta-cyclase' |
| NanoG1409 | 'lycopene beta-cyclase' |
| NanoG1414 | 'phytoene desaturase' |
| NanoG1423 | '2-c-methyl-d-erythritol 4-phosphate cytidylyltransferase' |
| NanoG1424 | 'geranylgeranyl diphosphate reductase' |
| NanoG1425 | '1-deoxy-d-xylulose-5-phosphate reductoisomerase' |
| NanoG1428 | '(E)-4-hydroxy-3-methylbut-2-enyl-diphosphate synthase' |
| NanoG1429 | 'geranylgeranyl diphosphate synthase' |
| NanoG1434 | '4-(cytidine 5''-diphospho)-2-c-methyl-d-erythritol kinase' |
| NanoG1435 | '2-c-methyl-d-erythritol 2,4-cyclodiphosphate synthase' |
| NanoG1437 | '(2e,6e)-farnesyl diphosphate synthase' |
| NanoG1439 | 'dimethylallyltranstransferase' |
| Tr_ADPHt_h | 'ADP:H+ symporter, chloroplast' |
| Tr_AMETt2h_c | 'S-Adenosyl-L-methionine reversible transport, chloroplast' |
| Tr_AMPt_h | 'AMP transport, chloroplast' |
| Tr_ASPt_h | 'Amino acid transporter (asp-L), chloroplast' |
| Tr_CMPt_h | 'CMP transport via diffusion, chloroplast' |
| Tr_CTPt_h | 'Plastid nucleotide transporter (CTP/ATP antiport)' |
| Tr_HISt_h | 'Amino acid transporter (his-L), chloroplast' |
| Tr_MG2t_h | 'Divalent cation (Mg2+) transport system, chloroplast' |
| Tr_UDPGALt_h | 'UDP-galactose:UMP antiporter, chloroplast' |
| Tr_UDPGt_h | 'UDP-glucose antiporter, chloroplast' |
| Tr_UDPt_h | 'UDP transport via diffusion, chloroplast' |
| B_adp[c] | 'B_adp[c]' |
| B_ahcys[c] | 'B_ahcys[c]' |
| B_amet[c] | 'B_amet[c]' |
| B_amp[c] | 'B_amp[c]' |
| B_asp-l[c] | 'B_asp-l[c]' |
| B_atp[c] | 'B_atp[c]' |
| B_cmp[c] | 'B_cmp[c]' |
| B_ctp[c] | 'B_ctp[c]' |
| B_his-l[c] | 'B_his-l[c]' |
| B_mg2[c] | 'B_mg2[c]' |

| | |
|---|---|
| B_udpgal[c] | 'B_udpgal[c]' |
| B_udpg[c] | 'B_udpg[c]' |
| B_udp[c] | 'B_udp[c]' |
| Ex_Photon | 'Photon import' |
| PSII_Photon | 'Directing photons to PSII' |
| PSI_Photon | 'Directing photons to PSI' |
| S0S1 | 'Manganese cluster S-cycling, S0 -> S1' |
| S1S2 | 'Manganese cluster S-cycling, S1 -> S2' |
| S2S3 | 'Manganese cluster S-cycling, S2 -> S3' |
| S3S4 | 'Manganese cluster S-cycling, S3 -> S4' |
| S4S0 | 'Manganese cluster S-cycling, S4 -> S0' |
| P680P | 'Electron transport P680 -> Pheophytin a' |
| YZP680 | 'Electron transport Yz -> P680' |
| PQA | 'Electron transport P680 -> QA' |
| QAQB1 | 'First reduction of QB' |
| QAQB2 | 'Second reduction of QB' |
| QBPQH2 | 'QB enters the PQ pool' |
| PQPSII | 'Docking of PQ to PSII -> QB(ox)' |
| PQH2R | 'Reduction of Rieske ISP in cyt b6f complex' |
| RISPCF | 'Electron transfer from Rieske ISP to Cyt f' |
| CFPC | 'Electron transfer from Cyt f to PC' |
| PQRHBP | 'Electron transfer from PQ radical to Heme bp' |
| HBPHBNHCN1 | 'First reduction of Heme bn/Heme cn by Heme bp' |
| HBPHBNHCN2 | 'Second reduction of Heme bn/Heme cn by Heme bp' |
| HBNHCNPQH2 | 'Re-oxidation of Heme bn/Heme cn and generation of secondary PQH2' |
| P700A0 | 'Excitation of P700' |
| PCP700 | 'Re-reduction of P700 by PC' |
| A0A1 | 'Electron transfer from A0 to A1' |
| A1FX | 'Electron transfer from A1 to Fx' |
| FXFB | 'Electron transfer from Fx to FB' |
| FBFA | 'Electron transfer from FB to FA' |
| FAFD | 'Electron transfer from FA to Ferredoxin' |
| FDFNR1 | 'First reduction of FNR' |
| FDFNR2 | 'Second reduction of FNR' |
| FNRNADPH | 'Reduction of NADP+ by FNR' |
| ATPSh | 'ATP synthase' |
| Tr_H2O_u | 'H2O transport, thykaloid' |
| Tr_O2_u | 'O2 transport, thykaloid' |
| ala-l[h] | 'L-alanine import to chloroplast' |

| | |
|---|---|
| B_ala-l[c] | 'B_ala-l[c]' |
| arg-l[h] | 'L-Arginine import to chloroplast' |
| B_arg-l[c] | 'B_arg-l[c]' |
| asn-l[h] | 'L-Asparagine chloroplast import' |
| B_asn-l[c] | 'B_asn-l[c]' |
| Tr_pro_h | 'L-Proline transport c <-> h' |
| B_pro-l[c] | 'B_pro-l[c]' |
| R04199_h | '2,3,4,5-tetrahydrodipicolinate:NADP+ 4-oxidoreductase' |
| R07613_h | 'LL-2,6-diaminoheptanedioate:2-oxoglutarate aminotransferase' |
| R02735_h | 'LL-2,6-Diaminoheptanedioate 2-epimerase' |
| R00451_h | 'meso-2,6-diaminoheptanedioate carboxy-lyase' |
| R00650_h | 'Homocysteine S-methyltransferase' |
| R00691_h | 'Arogenate dehydratase' |
| R00732_h | 'Arogenate dehydrogenase' |
| R01213_h | 'Isopropylmalate synthase' |
| R10170_h | 'Isopropylmalate isomerase' |
| R04426_h | 'Isopropylmalate dehydrogenase' |
| R01652_h | '2-Oxoisocaproate synthesis (spontaneous)' |
| R01090_h | 'Branched-chain aminotransferase' |
| @Chl_protprod | 'Chlamydomonas chloroplast protein production' |
| R06946_h | 'Zeaxanthin epoxidase' |
| R06947_h | 'Zeaxanthin epoxidase' |
| R06948_h | 'Neoxanthin synthase' |
| R01195_h | 'Ferredoxin:NADP+ oxidoreductase' |
| B_c182coa[c] | 'B_c182coa[c]' |
| c182coa_h | 'C18:2-CoA transport c -> h' |
| R02241_c182_h | 'Acyl-CoA:1-acyl-sn-glycerol-3-phosphate 2-O-acyltransferase' |
| B_c183coa[c] | 'B_c183coa[c]' |
| c183coa_h | 'C18:3-CoA transport c -> h' |
| R02241_c183_h | 'Acyl-CoA:1-acyl-sn-glycerol-3-phosphate 2-O-acyltransferase' |
| B_c204(6)coa[c] | 'B_c204(6)coa[c]' |
| c204(6)coa_h | 'C20:4(6)-CoA transport c -> h' |
| R02241_c204_h | 'Acyl-CoA:1-acyl-sn-glycerol-3-phosphate 2-O-acyltransferase' |
| B_c205(3)coa[c] | 'B_c205(3)coa[c]' |
| c205(3)coa_h | 'C20:5(3)-CoA transport c -> h' |
| R02241_c205_h | 'Acyl-CoA:1-acyl-sn-glycerol-3-phosphate 2-O-acyltransferase' |
| B_@Chl@Pha_dhacoa[c] | 'B_@C@P_dhacoa[c]' |
| @Chl@Pha_Tr_dhacoa[h] | 'DHA transport c -> h (Phaeo and Chlamy specific rx)' |
| @Chl_dhag3psynth | 'Glycerol-3-phosphate:acyl-coa acyltransferase C22:6 (Chlamy specific)' |

| | |
|---|---|
| @Chl_R02241_c226_h | 'Acyl-CoA:1-acyl-sn-glycerol-3-phosphate 2-O-acyltransferase (Chlamy specific)' |
| @Chl_dg226synth | 'Phosphatidate phosphatase (n-C22:6) (Chlamy specific)' |
| @Chl_mgdg226synth | 'UDPgalactose 1,2-diacylglycerol 3-beta-D-galactosyltransferase (n-C22:6) (Chlamy specific)' |
| @Chl_dgdg226synth | 'Galactolipid galactosyltransferase (Chlamy specific)' |
| @Chl_sqdg226synth | 'Sulfoquinovosyltransferase (n-C22:6) (Chlamy specific)' |
| mgdg180synth | 'UDPgalactose 1,2-diacylglycerol 3-beta-D-galactosyltransferase (n-C18)' |
| dgdg180synth | 'Galactolipid galactosyltransferase' |
| sqdg180synth | 'Sulfoquinovosyltransferase (n-C18)' |
| @Chl_mgdg183synth | 'UDPgalactose 1,2-diacylglycerol 3-beta-D-galactosyltransferase (n-C18 3)' |
| @Chl_dgdg183synth | 'Galactolipid galactosyltransferase' |
| @Chl_sqdg183synth | 'Sulfoquinovosyltransferase (n-C18 3)' |
| @Chl_mgdg_pf | 'Chlamydomonas MGDG pool formation' |
| @Chl_dgdg_pf | 'Chlamydomonas DGDG pool formation' |
| @Chl_sqdg_pf | 'Chlamydomonas SQDG pool formation' |
| @Chl_ccmemprod | 'Chlamydomonas chloroplast membrane lipid pool formation' |
| @Chl_R09067_h | 'Chlorophyllide-b:phytyl-diphosphate phytyltransferase' |
| @Chl_R07851_h | 'Alpha-cryptoxanthin,reduced ferredoxin [iron-sulfur] cluster:oxygen 3-oxidoreductase' |
| @Chl_pigm | 'Chlamydomonas pigment fraction assembly' |
| @Chl_bio | 'Chlamydomonas chloroplast biomass' |

Note: The list reactions is specifically *Chlamydomonas reinhardtii*-related, however the model uses *Nannochloropsis gaditana* as a base, hence all reactions common to both organisms have a "Nano" designation.
